# Umbilical Cord Blood Diagnostics for Early Onset Sepsis in Premature Infants: Detection of Bacterial DNA and Systemic Inflammatory Response

**DOI:** 10.1101/200337

**Authors:** Leena B. Mithal, Michael Malczynski, Chao Qi, Stefan Green, Patrick C. Seed, Karen K. Mestan

**Affiliations:** Pediatrics, Feinberg School of Medicine, Northwestern University; Clinical Microbiology Laboratory, Northwestern Memorial Hospital; Pathology, Northwestern University Feinberg School of Medicine; Center for Genomic Research, University of Illinois at Chicago

## Abstract

This work has been supported by a Thrasher Research Fund Early Career Award (#12573), National Institutes of Health (K23 HL093302 [KKM], R01GM108494 [PCS])), the Gerber Foundation, the Hartwall Foundation, and the Northwestern Memorial Foundation (Friends of Prentice Grants Initiative).

**Disclosure statement:** The authors of this manuscript have no conflicts of interest to disclose.

**Category of Study:** Translational

## Introduction

Sensitive and specific diagnosis of sepsis is difficult in newborn infants. The diagnosis of sepsis is particularly challenging in preterm infants who are disproportionately affected by invasive infection in the first days of life. Early onset sepsis (EOS) is commonly the result of an intrauterine infection^1-3^. Perinatal risk factors and clinical signs of EOS are common in premature infants even without culture-positive EOS, limiting their utility^4^. Diagnostic indecision leads to increased morbidity due to delayed treatment and overuse of empiric broad-spectrum antibiotics^5^. Blood culture, the current gold standard, is problematic: culture is slow and the sensitivity is limited due to maternal antibiotics, small specimen volumes, and low pathogen density per sampled blood volume^6,7^. Other postnatal laboratory tests such as white blood cell indices and C-reactive protein (CRP) have poor specificity^8-10^. Thus, in the absence of reliable diagnostics for EOS, infants are frequently treated for presumed, culture-negative sepsis (PS) with antimicrobial agents, the most commonly used class of medications in the NICU ^11^. Early life, prolonged antibiotic exposure may lead to adverse consequences including necrotizing enterocolitis, antibiotic resistance, fungal infections, and hearing loss^12-15^. Emerging data suggest these early life antibiotic exposures may also have longstanding effects on immune and metabolic physiology^16^. In sum, these data highlight the critical need for accurate diagnostic tools for EOS to foster targeted antibiotic use in premature infants.

Evaluation of umbilical cord blood (CB) may aid in diagnosis of neonatal pathology, including EOS^17-19^. CB most closely reflects the inflammatory state of the intrauterine environment, where EOS originates in most preterm births. We have recently shown that three acute phase reactant (APR) proteins: serum amyloid A (SAA), CRP, and haptoglobin (Hp) are significantly elevated in CB of premature infants with EOS. These APRs have the potential to distinguish preterm infants with and without true infection at the time of birth^20^. We predicted that molecular detection of pathogens in CB would further enhance CB host response biomarkers in the diagnosis and exclusion of EOS.

CB 16S rDNA PCR has the potential advantage of earlier, more accurate bacterial detection for EOS diagnosis compared with postnatal blood culture. At this time, the utility of 16S rDNA PCR of CB as a culture-independent method of bacterial detection to diagnose EOS remains unknown. The objective of this study was to compare CB 16S rDNA PCR followed by Sanger amplicon sequencing (PCR/Sanger-Seq) from preterm infants with confirmed sepsis (cEOS), PS, late onset sepsis (LOS) and no sepsis (controls). We aimed to correlate postnatal blood culture results and CB inflammatory markers (SAA, CRP, Hp) with PCR/Sanger-Seq of CB for detection of bacterial pathogens. We hypothesized that CB PCR/Sanger-Seq would identify bacteria in the setting of cEOS and help distinguish PS infants with true infection from a subset less likely to have infection. We further sought to explore the contribution of next generation sequencing (NGS) of CB 16S rDNA amplicons to the diagnosis of EOS in a subset of infants.

## Methods

### Study design

We conducted a nested case-control study of premature infants (<37 weeks gestational age) enrolled in a longitudinal birth cohort at Prentice Women’s Hospital (Chicago, IL). Enrollment was dependent on parental consent and availability of CB. At the time of study, pertinent clinical data were available for 1,100 enrolled infants born between 2008 and 2014. Per the parent study protocol, umbilical venous CB was collected at delivery by sterile venipuncture into EDTA tubes, spun at 3,000rpm for 10 minutes in a refrigerated tabletop centrifuge. Plasma was immediately separated from the red blood cell (RBC) layer and buffy coat. All aliquots were immediately stored at −80°C, de-identified and coded. Comprehensive clinical data of maternal and infant variables were collected, including demographics, maternal risk factors, indication for preterm delivery, laboratory values, culture results, and administered antibiotics. Placental histopathology results were available, including maternal and fetal acute inflammation variables as previously described^20,21^.

### Patients

Based on culture results, antibiotic treatment and laboratory data, patients were classified in the following categories on the basis of proven or presumed sepsis: cEOS, PS, LOS, and controls, according to pre-determined criteria based upon National Institute of Child Health and Human Development definitions for premature infants and as described in our previous work (Figure 1)^20^. Infants were considered to have cEOS if a positive blood culture was present in the first 72 hours of life (HOL) for which they received an antibiotic course (≥5 days). Patients with questionable true EOS pathogens, such as *Corynebacterium spp.*, or with viral or fungal sepsis were excluded. PS infants had negative blood cultures and ≥2 abnormal laboratory results from peripheral blood in the first 72 HOL. PS infants received an antibiotic course which began within 72 hours after birth. Abnormal postnatal laboratory results were: CRP ≥1mg/dL^22^, absolute neutrophil count (ANC) outside normal range for GA^23^, and immature-to-total (I:T) neutrophil ratio >0.2^24^. LOS patients had positive blood culture after 72 HOL treated with an antibiotic course and no EOS (proven or presumed). Control patients had no positive culture throughout hospitalization and received no more than 4 days of antibiotics (Table 1). The majority of control patients (79.7%) received 2-3 days of antibiotics as “rule-out” of sepsis pending negative culture result. PS, LOS, and control patients were frequency matched based on GA and birth weight to cEOS cases. CB APRs were measured by magnetic bead immunoassays (Bio-Rad Laboratories Inc.; Hercules, CA), and placental histopathologic inflammation was characterized as previously described^20^. An automated continuous-monitoring blood culture system, BacT/ALERT (bioMerieux, Durham, NC) is used in the microbiology laboratory in Northwestern Medicine Hospitals as standard patient care. These results were retrospectively obtained by review of electronic medical records.

**Figure 1.**
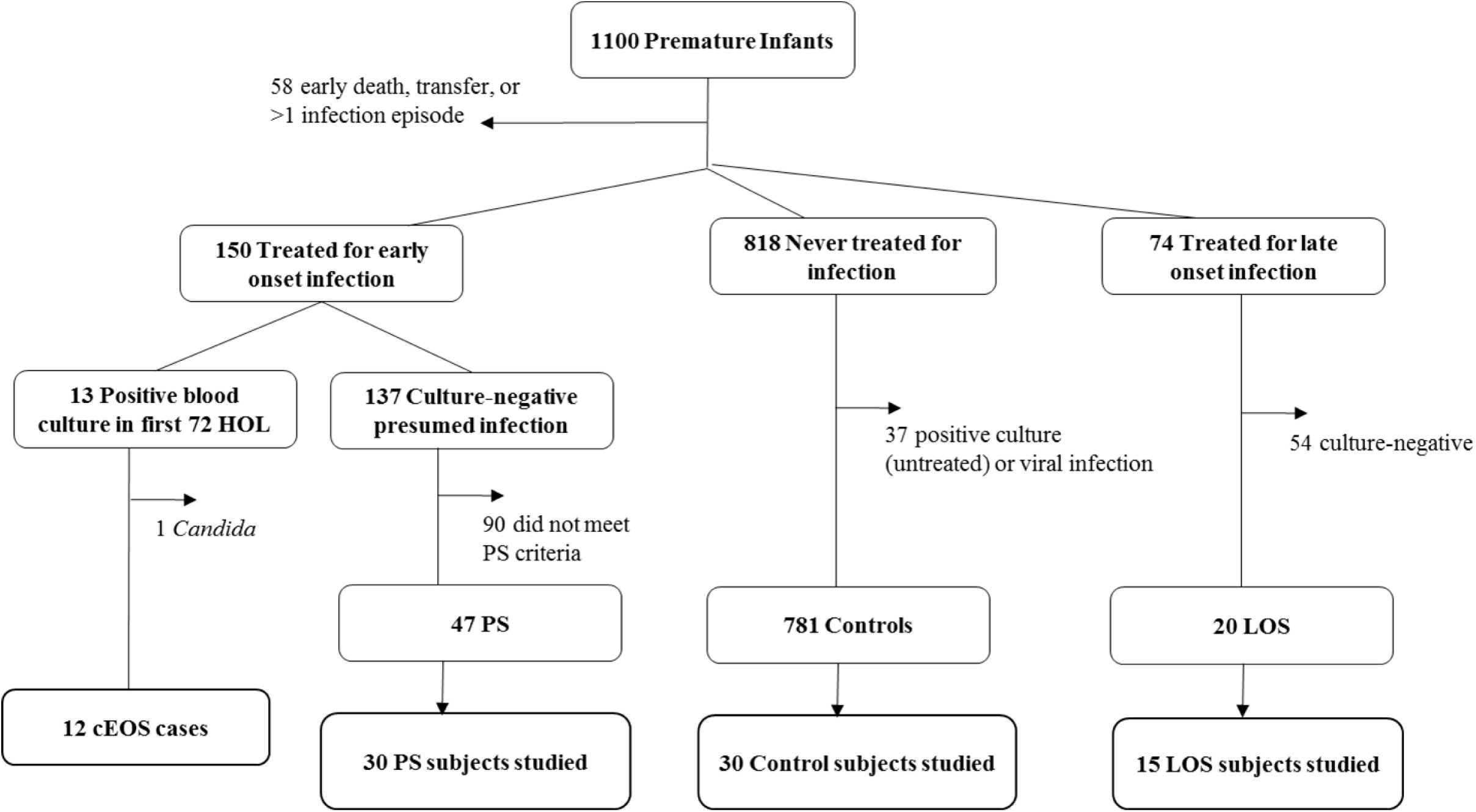
Study design and patient cohort. Of 1,100 enrolled preterm infants, 12 were confirmed early onset sepsis (cEOS: positive blood culture with bacterial pathogen <72 HOL and antibiotic treatment ≥5 days). Presumed sepsis (PS) subjects received antibiotic treatment course initiated within 72 HOL, had no positive sterile site culture, and had ≥2 abnormal laboratory criteria (↓ANC, ↑ I:T ratio, ↑CRP). Control patients had no positive culture or antibiotic course >4 days. Late onset sepsis (LOS) infants had a positive blood culture at >72 HOL treated with antibiotic course. Patients with both EOS and LOS were excluded. PS, LOS and control patients were frequency matched to cEOS patients by gestational age (GA) and birth weight (BW).

**Table 1.** Case definitions and demographics

| Variable | Control<br>n=30 | cEOS<br>n=12 | PS<br>n=30 | LOS<br>n=15 | Total Sample<br>N=87 |
| --- | --- | --- | --- | --- | --- |
| **Positive blood culture** | No | Yes, <72 HOL | No | Yes, >72 HOL | - |
| **Antibiotic treatment course** | No | Yes | Yes | Yes | - |
| **Gestational age, weeks**<br>median (IQR) | 29.9 (27.7-32.3) | 30.2 (27.4-32.8) | 29.6 (28.1-32.0) | 28.3 (26.1-29.3) | 29.4 (27.6-31.9) |
| **Birthweight, grams** | 1365<br>(1055-1770) | 1465<br>(1003-2051) | 1360<br>(1000-1665) | 1075*<br>(875-1155) | 1230<br>(995-1670) |
| **Male gender, N(%)** | 12 (40) | 8 (67) | 24 (80)* | 9 (60) | 53 (61) |
| **Multiple gestation** | 11 (37) | 3 (25) | 13 (43) | 5 (33) | 32 (37) |
| **Vaginal delivery** | 15 (50) | 9 (75) | 11 (37) | 7 (47) | 42 (48) |
| **Prolonged rupture of membranes** | 8 (27) | 8 (67) | 11 (37) | 4 (27) | 27 (36) |
| **Preeclampsia** | 6 (20) | 0 | 7 (23) | 3 (20) | 16 (18) |
| **Clinical chorioamnionitis** | 0 | 5 (42)* | 4 (13) | 0 | 9 (10) |
| **Placental histopathologic AI<sup>b</sup>** | 17 (57) | 12 (100)<sup>a</sup> | 9 (32) | 5(37) | 43 (49) |

Bonferroni adjusted p-values of 0.01 per test were used to determine statistical significance.
\* p<0.01 versus control group
<sup>a</sup> p<0.01 versus control, PS, and LOS groups
<sup>b</sup> AI=acute inflammation, (maternal or fetal AI as previously described)<sup>21</sup>. Placental data not available for 2 PS patients (n=28)

### 16S rDNA PCR amplification and Sanger sequencing (PCR/Sanger-Seq)

Archived de-identified coded CB samples were identified and obtained from the parent study. DNA extraction and 16S rDNA PCR were performed in the clinical microbiology laboratory at Northwestern Memorial Hospital on each sample using established protocols. Briefly, DNA was extracted from the CB buffy coat with the QIAamp DNA Mini Kit (Qiagen Sciences, LLC, Louisville, KY). For each batch of extractions, a negative extraction control was performed using all reagents minus blood. Extracted blood DNA was subjected to PCR amplification of 16S rDNA by using the following V3-V4 universal primer pair: 8F: 5’-GAGAGTTTGATYMTGGCTCAGRRYGAACGCT-3’ and 806R:

5’GGACTACCAGGGTATCTAAT-3’. The primers provide a PCR product of 806 bp. Each PCR reaction (50μl) consisted of 10x Buffer, Taq polymerase (Invitrogen Platinum taq DNA polymerase high fidelity), 4mM MgSO_4_, dNTP, DNAse/RNAse-free H_2_0, and 4μl DNA template. Cycling conditions included a 5 minute denaturing step at 95C followed by 34 cycles of 30 seconds at 94C, 30 seconds at 55C, and 60 seconds at 68C. Sterile water and *K pneumoniae* ATCC® BAA-1705 were used as negative and positive DNA controls, respectively^25^. The PCR results were considered positive when visible PCR products of the correct size were found on electrophoresis gel. PCR results were considered negative when visible PCR products of the correct size were not visualized following ethidium bromide staining and UV illumination. In circumstances where the negative extraction controls produced a clear moderate band, the accompanying batch of PCR analysis was disgarded and repeated. In cases where possible faint amplicons were visualized from the negative control, the bands were purified (QIAquick PCR Purification Kit, Qiagen) and sequenced in the NUSeq Core Facility of Northwestern University. The results were poor sequence output or sequences corresponding to *Cloacibacterium*. In the clinical lab, if a patient sample is less visible than a faint control band, the result is considered negative. The Sanger sequencing was carried out by using BigDye(tm) Terminator v3.1 Cycle Sequencing Kit. The reaction was performed on a GeneAmp® PCR System 9700 (ThermoFisher, Waltham, MA). After cleanup, the reaction product was loaded onto a 3730 DNA analyzer (ThermoFisher) and sequencing data was produced by Sequencing Analysis Software 6 (ThermoFisher). If there was very low or no sequence output (trimmed length <300bp), the result was considered negative. National Center for Biotechnology Information (NCBI) database BLAST was used for species identification.

### Next generation sequencing

In a subset of patients with sufficient remaining high-quality genomic DNA, high-throughput sequencing of 16S rDNA amplicons was performed (8 cEOS, 12 PS, 20 controls) through the University of Illinois at Chicago Core Genomics Facility. V3/V4 or V4 region 16S rDNA targeted primers were employed (341F: 5’-CCCTACGGGAGGCAGCAGCTACGGGAGGCAGCAGCCTACGGGAGGCAGCAG-3’or 515F: 5’-GTGCCAGCMGCCGCGGTAA-3’ and 806R: 5’-GGACTACHVGGGTWTCTAAT-3’ as previously described^26^. Amplicon barcoding was achieved using a second PCR, incorporating Illumina adapters (Fluidigm AccessArray barcoding system) and a sample specific multiplex identifier (MID) sequence. Barcoded sample amplicons were pooled and sequenced using an Illumina MiSeq sequencer (V2; 2x250 paired-end reads; Illumina Inc, San Diego, CA). We used 6 extraction blanks as methodologic controls: 2 with sterile water, 2 sterile water through extraction column, and 2 sterile water through extraction column using all reagents in the QIAamp DNA Mini Kit.

### Statistical analysis

Clinical variables were reported as medians (25th-75th interquartile range; IQR) or frequencies and percentages. The Kruskal-Wallis tests and chi-square or Fisher’s exact tests were used to identify differences in variables across sepsis categories as appropriate. For variables showing statistical significance, comparisons of sepsis categories were assessed using the Wilcoxon rank sum test and chi-square/Fisher’s exact tests. Bonferroni-adjusted p-values of p <0.01 were used to account for multiple comparisons (5 individual comparisons between sepsis groups). STATA® software, version 13.1 (College Station, TX) was used for this analysis.

Bioinformatics was using QIIME^27^ and the following packages in the R software environment: phyloseq (version 1.16.2), vegan (version 2.4-1), metagenomeSeq (version 1.14.2), reshape2 (version 1.4.2), ggplot2 (version 2.2.1), plyr (version 1.8.4), dplyr (version 0.5.0), biom (version 0.3.12), gss (version 2.1-6), ggbeeswarm (version 0.5.3), ggsignif (version 0.3.0), and Biostrings (version 2.40.2) REFS. Paired-end reads were merged and primer sequences removed. Low quality sequences were removed. Sequences were annotated using a custom QIIME pipeline, and biological observation matrices (BIOMs) were generated at taxonomic levels from phylum to species using the Greengenes 13_8 reference database ^28,29^. BIOMs were processed for α- and β- diversity (Bray-Curtis dissimilarity) analyses in the software package ’vegan’ (Community ecology package 10, 2007), implemented in the “R” software environment. For alpha-diversity analysis, Shannon Index values were generated for the genus level and Kruskal-Wallis test was used to for test significant difference across groups. Pairwise comparisons between SI of individual sepsis groups employed Wilcoxon tests. Beta-diversity analyses were performed by constructing nonmetric multi-dimensional scaling (NMDS) plots. Community similarity was tested using a pairwise adonis with Benjamini-Hochberg correction of microbial community composition.

### Ethics statement

The study was approved by the Northwestern University Feinberg School of Medicine Institutional Review Board. Parental written informed consent was obtained for each patient prior to enrollment. Subsequent investigation was conducted according to the principles expressed in the Declaration of Helsinki.

## Results

### Clinical data

Of 1,100 enrolled infants (Figure 1), there were 12 cases of cEOS from bacteria due to the following organisms: *Escherichia coli* (n=7), *Streptococcus agalactiae* (n=2), *Proteus mirabilis* (n=1), *Haemophilus influenzae* (n=1) and *Listeria monocytogenes* (n=1). Approximately 1% of infants had a blood-culture proven EOS, consistent with national reports, while 19% received a prolonged empiric antibiotic course for EOS^2^. The 87 patients presented here (cEOS, PS, LOS, controls) had a median gestational age of 29.4 weeks (IQR: 27.6-31.9) and median birth weight of 1230g (IQR: 995-1670). Demographic characteristics were similar across groups with no significant differences in gestational age, multiple gestation, route of delivery, prolonged rupture of membranes (PROM), and preeclampsia between groups (Table 1). The LOS group was lower birth weight than controls (p=0.01). Clinical chorioamnionitis was more prevalent in the cEOS group than LOS (p=0.01) or controls (p=0.001). The presumed sepsis group had a higher proportion of male gender than controls (p= 0.002). Placental histopathologic acute inflammation was higher in the cEOS group than each of the other categories (p<0.01).

### Sanger sequencing of 16S rDNA amplicons

All 12 cEOS patient CB produced 16S rDNA amplicon bands visualized by gelelectrophoresis. Sanger sequencing yielded species level identification for 9 of 12 patients. Three PCR/Sanger-Seq provided poor sequence homology/no species level identification from NCBI BLAST. In 8 subjects, PCR/Sanger-Seq identified the same organism as was cultured in postnatal blood (Table 2). In one infant with *E coli* in postnatal blood culture, cord blood Sanger sequencing identified *G vaginalis*. This infant was born by vaginal delivery without PROM or clinical chorioamnionitis, but did have stage 2 fetal acute inflammation, funisitis, and chronic inflammation on placental histopathology. Of the 30 PS infants, 21 had a negative PCR/SangerSeq result, 8 had a single species identified, and 1 had a band with 400bp of trimmed length sequence and no species level identification. As shown in Table 2, positive organisms in PS were: *S agalactiae* (n=2), *E coli, Enterobacter aerogenes, Haemophilus parainfluenzae, Streptococcus gallolyticus, Sneathia sanguineges,* and *Actinomyces neuii*. One of 15 LOS infants had positive band identified as *Sneathia amnii*, while the other 14 had negative PCR. Of the 30 control infants, 23 PCR/Sanger-Seq were negative while 7 had a single bacterial species identified by sequencing: *Lactobacillus crispatus* (n=3), *Mycoplasma hominis* (n=2), *Lactobacillus iners* (n=1), and *Prevotella bivia* (n=1). Using true positive and negative EOS patients (cEOS, LOS+controls) and considering a positive PCR/Sanger-Seq result as species level identification, the sensitivity of CB 16S rDNA PCR/Sanger-Seq for cEOS was 75%, specificity was 82.2%, and positive and negative predictive values were 53% and 92.5% respectively.

**Table 2.**
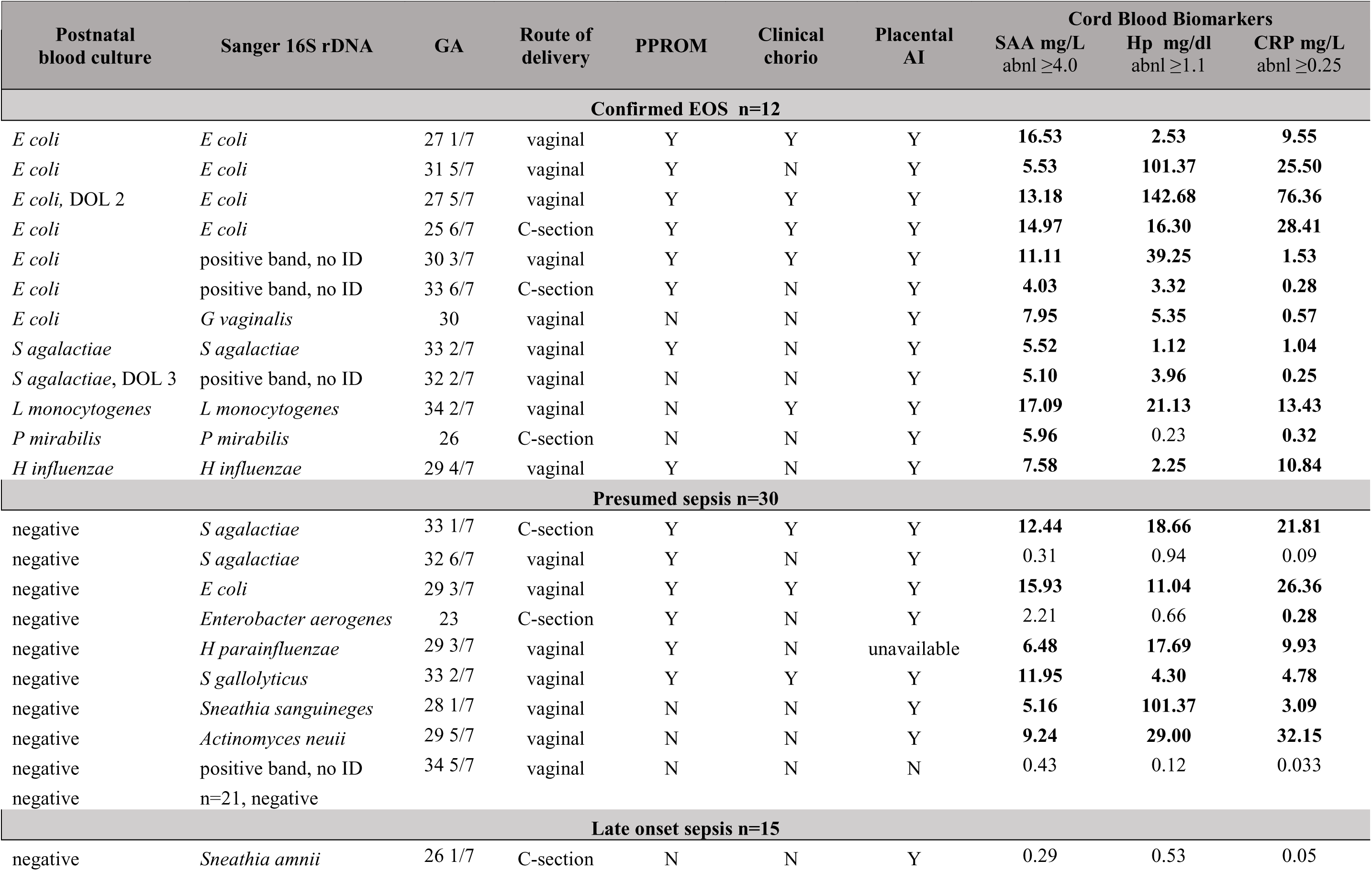
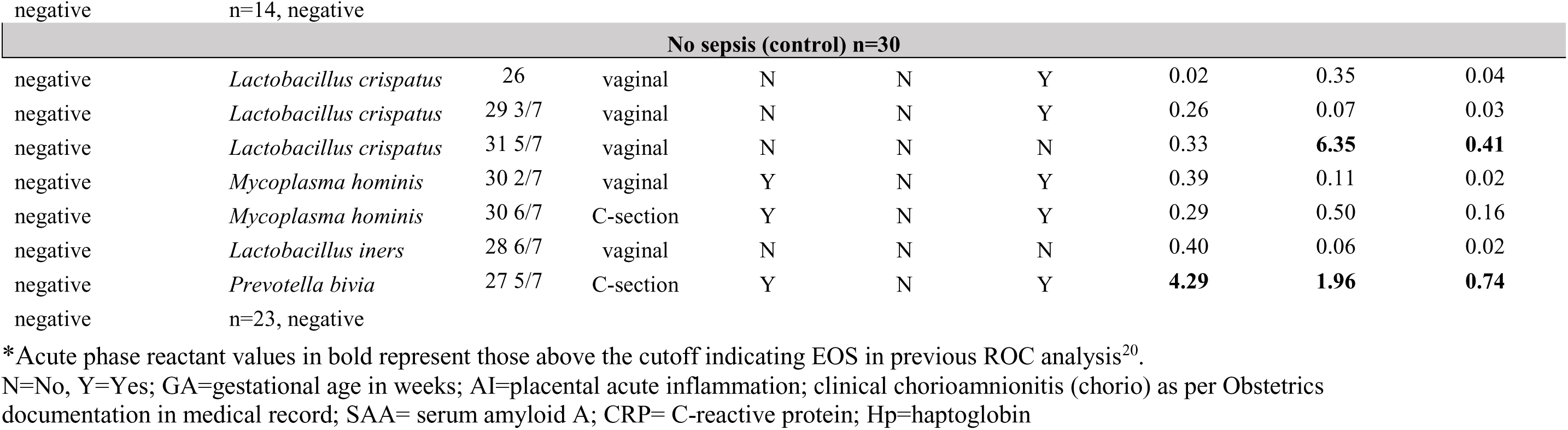
Cord blood Sanger 16S rDNA sequencing and acute phase reactants of sepsis groups

### Correlation between Sanger sequencing and cord blood acute phase reactants

We previously reported CB APR and placental histopathology results on this cohort^20^. SAA, CRP and Hp levels were significantly higher in CB of cEOS patients and a subset of PS patients compared to controls and LOS (Table 2; complete individual infant data in Supplemental Table 1). As shown in Figure 2, cEOS infants all had positive PCR/Sanger-Seq and elevated markers. In contrast, the control infants with positive PCR/Sanger-Seq as a whole did not have elevated markers, except 1 infant with Sanger identification of *Prevotella* and an elevation of all 3 CB APRs. This clinical picture was significant for 27 5/7 weeks GA, maternal antibiotics, premature preterm PROM, stage 2 maternal and fetal placental acute inflammation, neutropenia at birth, and treatment with 48 hours of empiric antibiotics to “rule-out” infection. The LOS infant with *Sneathia* and those with negative PCR/Sanger-Seq did not have elevated CB APRs.

**Figure 2:**
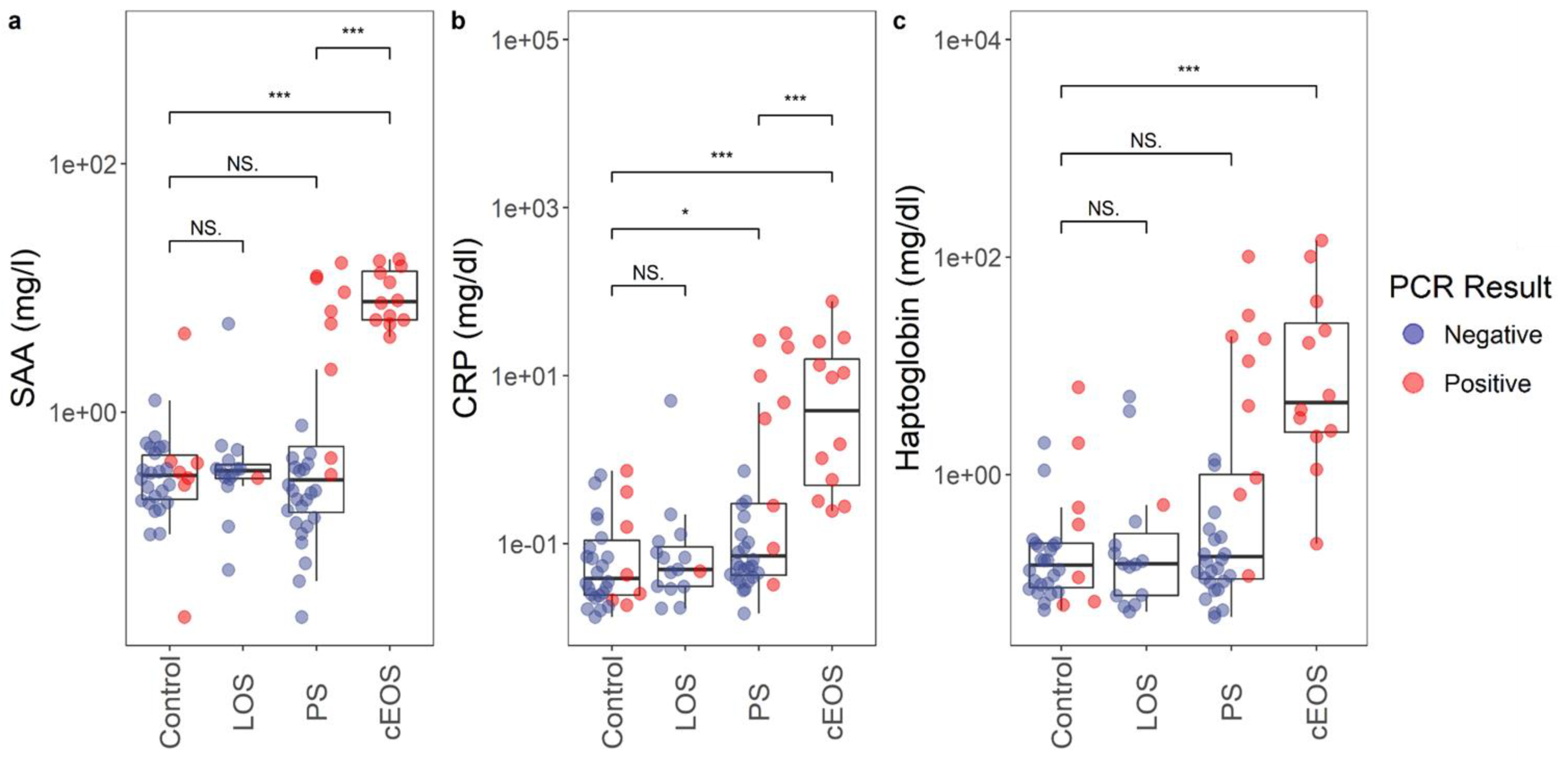
Cord blood 16S rDNA sequencing and acute phase reactants. Box and whisker plots displaying distribution biomarkers a) serum amyloid A [SAA], b) C-reactive protein [CRP], and c) haptoglobin [Hp] in each sepsis category (log-transformed). Red = positive 16S rDNA PCR/sequencing; blue = negative 16S rDNA PCR/sequencing result. Positive PCR in cEOS and PS groups were highly correlated with CB biomarker elevation.

Seven out of 8 PS subjects with positive CB PCR/Sanger-Seq had an elevated CB APR, based on previously established ROC cutoffs (analysis of cEOS and control infants)^20^. The one PS infant with normal CB APRs had PCR/Sanger-Seq identification of *S agalactiae*. Six of the 8 infants with positive Sanger had elevation of all 3 CB APRs. The PS subject with positive 16S rDNA band but with no species identification and all the PCR/Sanger-Seq negative PS infants did not have elevated CB APRs.

### Next generation sequencing

NGS results were obtained for 40 infants of 87 infants (8 cEOS, 12 PS, 20 controls) and 6 blanks. NGS detected bacterial DNA amplification in all CB samples. In 7 of 8 the cEOS samples, NGS demonstrated the taxa (typically genus level identification) of the pathogenic bacteria in postnatal blood culture, with variable percentage abundance (Figure 3a). In 1 cEOS sample, two organisms, *Escherichia* and *Gardnerella*, were identified by culture and PCR/Sanger-Seq, respectively. Less than one percent of the NGS abundance in that sample accounted for *Gardnerella,* while *Escherichia* was not identified. Each of the three PS subjects with positive PCR/Sanger-Seq had correspondingly high percentages of the concordant organism by NGS. The control infants with bacterial species identification on Sanger sequencing had the bacterial genus found on NGS, again with variable percent abundance.

**Figure 3.**
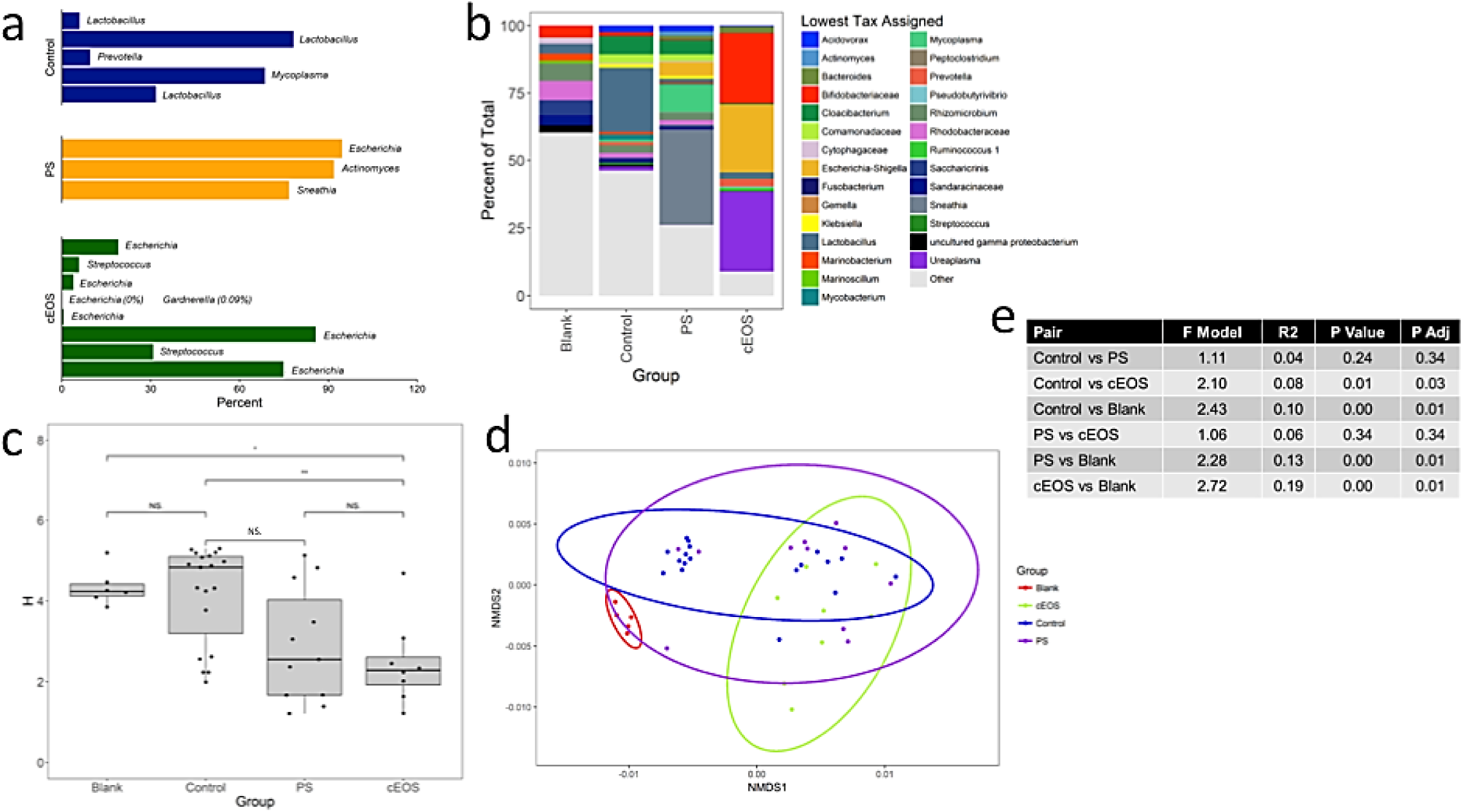
Next generation sequencing of 16S rDNA amplicons from cord blood. a) Percent abundance of bacterial species of interest (species in postnatal blood culture or 16S rDNA Sanger sequencing result in PS and control infants). b) Relative abundance of bacterial taxa in aggregate for each group: blanks, control, PS, cEOS. c) Alpha-diversity analysis. *Pairwise comparison of SI between cEOS and controls is significant (p=0.03) d, e) Beta-diversity analysis. Nonmetric multi-dimensional scaling (NMDS) plot constructed using Bray-Curtis dissimilarity between microbial community composition of samples at genus level: 46 samples (8 cEOS, 12 PS, 20 controls, 6 blanks). Ellipses show 95% confidence interval of each group. In pairwise adonis comparisons, blanks as a group were significantly different than each other group (adjusted p=0.01), cEOS and controls were different (adjusted p=0.03).

The relative abundance of the top 10 bacterial taxa per sample aggregated by clinical group are shown in Figure 3b. Among the cEOS patient samples Bifidobacteriaceae (which includes *Gardnerella spp.*), *Ureaplasma,* and *Escherichia-Shigella* predominated. PS samples also had high *Escherichia-Shigella* content but otherwise demonstrated a different aggregated composition. Control infants uniquely had a high percentage of *Lactobacillus*. The blanks did have bacterial DNA amplified by NGS, but was most diverse with no predominant organism/taxa in any sample.

The predominance of particular bacteria and the differences in bacterial community makeup is demonstrated in the alpha-diversity analysis (Figure 3c). The Shannon Index value was not significant across all groups (Kruskal-Wallis test, p=0.47). However pairwise comparisons of Shannon Index between cEOS and controls was also significantly different, with cEOS being less diverse (Wilcoxon test, p=0.003). The beta-diversity is displayed in Figures 3d and 3e. The NMDS plot shows ellipses at 95% confidence interval for each group and demonstrates a clear segregation of the “blank” samples. Two major groupings otherwise are apparent. One group has more than half of the control samples with several PS samples. The other more dispersed group has Control, PS, and the cEOS samples. In pairwise adonis comparisons, blanks as a group were significantly different than each other group (adjusted p=0.01), cEOS and controls were different (adjusted p=0.03), and PS was not significantly different than cEOS or controls (adjusted p=0.34 each).

## Discussion

This study reports and compares pathogen identication between cord blood PCR with Sanger-based 16S rDNA sequencing, next generation sequencing of 16S rDNA amplicons from cord blood, cord blood APR levels, and postnatal venous blood culture in preterm infants with confirmed, presumed, and no EOS. We found that amplification and sequencing of 16S rDNA can identify pathogenic bacteria in cases of EOS, including in some cases of culture-negative presumed sepsis, and if negative, may aid in ruling-out an infection. Rapid, molecular-based testing of CB for early pathogen detection in EOS could address an important clinical challenge.

Our results suggest that CB PCR/Sanger-Seq could provide organism-specific information helpful for earlier diagnosis of EOS, correctly identifying the implicated bacteria in two-thirds of cEOS cases and with a high NPV to exclude EOS in uninfected controls. Furthermore, pathogenic bacteria were detected in a number of PS patients with negative postnatal blood culture. Identification of a particular pathogen could allow tailored antibiotic therapy, rather than empiric broad spectrum antibiotics. Molecular detection also can identify fastidious organisms that are difficult to isolate in culture and has the potential to identify novel pathogens, such as *Sneathia* and *Actinomyces*. CB APR data aligns with our CB 16S rDNA results, suggesting that PS infants with negative CB PCR and no APR elevation may not have had true infection. Thus, the NPV of CB markers are reinforced by the data combining two approaches: detection of pathogen and host inflammatory response. With further validation, these CB diagnostics have the potential to significantly reduce antibiotic exposure for preterm infants perhaps incorrectly treated with prolonged antibiotics for possible sepsis in the absence of a positive pathogen identification (approximately 70% of our PS group). 16S rDNA PCR with Sanger sequencing is becoming more widely available in hospital clinical microbiology laboratories for sterile site fluids, and its use as an adjunct EOS diagnostic is feasible, with some advantages relative to the blood culture gold standard.

Previous studies on *postnatal* blood 16S rDNA assays for sepsis in the NICU have had design limitations, combining clinical and culture-proven sepsis across a broad timeline that includes both EOS and LOS. This body of evidence suggests improved sensitivity of 16S rDNA in detection of a bacterial pathogen both alone and in combination with microbial culture, but not yet sufficient to replace culture^30,31^. To our knowledge, the only recently published data on CB 16S rDNA sequencing has been by Wang and colleagues who investigated the link between intraamniotic infection and EOS through paired amniotic fluid and cord blood 16S rDNA PCR^32^.

In cases of EOS, the same bacterial pathogen was confirmed to be present in both amniotic fluid and cord blood. Similar to our PS group, CB PCR identified pathogens in some infants with presumed sepsis and negative culture, in the setting of intraamniotic infection. In their controls, however, no bacterial DNA was detected in either CB or amniotic fluid. In our study, CB of 6 of 30 control infants had *Lactobacillus* or *Mycoplasma spp* DNA by Sanger sequencing, all with normal CB APRs. These genera were not identified in the extensive negative controls. The significance of this bacterial DNA remains unclear, although we know that these infants did not receive antibiotic therapy effective against atypical bacteria and did well clinically. A previous study by Goldenberg *et al* found associations between the presence of certain bacteria in cord blood culture (*U. urealyticum, M. hominis*) and both preterm birth and adverse neonatal outcomes^33^. They also reported increased CB IL-6 and systemic inflammatory response syndrome in infants with positive CB culture with these organisms. In contrast, our data showed no consistent association between positive CB PCR and placental inflammation or infant inflammatory response among the control infant samples. Of the 6 control infants with positive PCR/Sanger-Seq, placental inflammation was present in 4 of 6, fetal acute inflammation in 2 of 6, and only 1 had elevated CB APRs. These rates were not different from the control infants with negative CB 16S rDNA PCR. Our results are more consistent with a 2005 study by Jimenez *et al*, in which 16S rDNA sequencing identified bacterial DNA (*E faecium, S epidermidis, S sanguinis, P acnes*) from CB of healthy full term infants delivered by Caesarian route. In sum, our data combining 16S rDNA PCR with CB inflammatory markers support a model whereby *in utero* microbial exposure occurs even in the absence of a systemic inflammatory response.

Because direct Sanger sequencing (without cloning) can be negatively impacted when elevated microbial diversity is present, we explored NGS of cEOS, PS, and control infants to characterize prenatal microbial DNA exposures around the time of birth. Our results demonstrate the presence of diverse microbial DNA in CB of preterm infants, with and without infection, and in infants with Caesarean and vaginal delivery alike. CB of 3 cEOS infants had a positive 16S rDNA band that did not produce valid Sanger sequencing. NGS demonstrated genomic presence of the postnatal pathogen in CB of these infants, though in lower percentages and also detected other organisms typically considered to be intraamniotic infection pathogens in high abundance (such as *Ureaplasma*, *Gardnerella,* and *Mycoplasma*). These data suggest that the fetus may have diverse microbial exposure *in utero* in the setting of infection. We hypothesis that a true pathogen within the setting of exposure to specific but diverse microbes may contribute to the development of an invasive infection and the associated inflammatory response. Alpha-diversity was significantly higher in control versus cEOS infants. Low diversity may be an indication of a shift from typical, diverse, and non-inflammatory microbial exposure to an *intrauterine* exposome with a bacterial makeup that promotes development of EOS and, in some cases, promotes emergence of a dominant bacterial pathogen. PS patients had some clinical signs and postnatal laboratory results concerning for sepsis, which prompted evaluation and empiric therapy despite negative cultures. Although our CB APR data were not elevated in a majority of these infants with negative Sanger sequencing, perhaps there are particular bacterial DNA exposures that lead to an inflammatory phenotype in the hours after delivery, thus evading detection through CB analysis. The concept of bacterial DNA triggering immunologic response in infant has been proposed, for instance in the case of TLR activation as reported by El Kebir and colleagues^34^. Further investigation of CB NGS may help us characterize the microbial exposures present in fetal circulation and understand microbe-microbe and microbe-host interactions in fetal development, preterm birth, EOS and the poorly understood culture-negative PS entity.

Despite a large enrollment cohort, this study is limited by the small sample size of groups, in particular cEOS, which reflects the real-life situation in the NICU where the number of culture-positive cases are few. For this reason, the duration of antibiotic treatment and clinical criteria are used to guide evaluation and treatment of infection, represented by our PS group. Given this reality, the strict sepsis categories and independent analysis of cEOS and PS groups is a strength of the study that allows the heterogenous PS group to be better defined, namely as infants with and without elevation in APRs and positive PCR/Sanger-Seq. NGS is a particularly high sensitivity technique with potential for contamination. Sterile CB collection and laboratory technique with multiple positive and negative PCR controls were used to minimize contamination as a confounder. While blood culture obtained within the first 48 hours of life most closely reflects circulating pathogens at birth, validation with cultures from the CB sample may have provided additional information about the timing of bacterial transmission and/or possibility of contamination. Lastly, while most all births in this study involved maternal antibiotic administration, differences in timing, dose, and type of antibiotic given to the mother was beyond the scope of this study but needs careful consideration in future studies. Finally, 16S rDNA NGS is limited by the inability for species level identification, detection ofviralal and microeukaryotic organisms

Our results of CB 16S rDNA PCR/sequencing combined with precise subject sepsis categorization and CB APR data provide an important foundation for further investigation of 1) molecular methods for both diagnosis of EOS and pathogen detection and 2) the *in utero* microbial DNA exposures that may shape an infant’s prenatal and postnatal course. In conclusion, it appears that CB contains evidence of a primary EOS pathogen in the majority of cases of EOS in preterm infants and can identify bacteria when a postnatal culture may not. 16S rDNA PCR detection has potential to effectively rule-out infection, particularly in combination with CB APRs. Typically commensal vaginal organisms have been identified in preterm infants without infection and, in some cases, are overrepresented in the setting of EOS with a different classic pathogen. These findings need to be verified by further investigation of fresh samples, in a larger, prospective cohort, and with sophisticated analysis of complex microbial exposures evidenced by deep sequencing. Such studies would provide the data required to incorporate CB pathogen detection into standard practice and may improve the understanding and management of EOS in the premature infant population.

## Acknowledgements

We would like to acknowledge the technical support of the NUSeq Core Facility of Northwestern University, the University of Illinois at Chicago Core Genomics Facility, the Northwestern Biostatistics Collaboration Core, Northwestern Enterprise Data Warehouse (EDW) and the Comprehensive Metabolic Core of Northwestern University. We thank Linda Ernst, MD for her expertise in interpretation and coding of placental histopathology and Lucy Minturn, MA and Juanita Saqibuddin, RN, of the Prentice NICU Cord Blood Study. We acknowledge our funding sources: the National Institutes of Health (K23HL093302 [KM], R01GM108494 [PCS]), Friends of Prentice Grants Initiative, the Gerber Foundation, The Hartwell Foundation, and Thrasher Research Fund. Finally, we acknowledge the children and their parents who participate in the longitudinal cohort study at Prentice Women’s Hospital.

